# A tandem-repeat dimeric RBD protein-based COVID-19 vaccine ZF2001 protects mice and nonhuman primates

**DOI:** 10.1101/2021.03.11.434928

**Authors:** Yaling An, Shihua Li, Xiyue Jin, Jian-bao Han, Kun Xu, Senyu Xu, Yuxuan Han, Chuanyu Liu, Tianyi Zheng, Mei Liu, Mi Yang, Tian-zhang Song, Baoying Huang, Li Zhao, Wen Wang, A Ruhan, Yingjie Cheng, Changwei Wu, Enqi Huang, Shilong Yang, Gary Wong, Yuhai Bi, Changwen Ke, Wenjie Tan, Jinghua Yan, Yong-tang Zheng, Lianpan Dai, George F. Gao

## Abstract

A safe, efficacious and deployable vaccine is urgently needed to control COVID-19 pandemic. We report here the preclinical development of a COVID-19 vaccine candidate, ZF2001, which contains tandem-repeat dimeric receptor-binding domain (RBD) protein with alum-based adjuvant. We assessed vaccine immunogenicity and efficacy in both mice and non-human primates (NHPs). ZF2001 induced high levels of RBD-binding and SARS-CoV-2 neutralizing antibody in both mice and NHPs, and also elicited balanced T_H_1/T_H_2 cellular responses in NHPs. Two doses of ZF2001 protected Ad-hACE2-transduced mice against SARS-CoV-2 infection, as detected by reduced viral RNA and relieved lung injuries. In NHPs, vaccination of either 25 μg or 50 μg ZF2001 prevented infection with SARS-CoV-2 in lung, trachea and bronchi, with milder lung lesions. No evidence of disease enhancement is observed in both models. ZF2001 is being evaluated in the ongoing international multi-center Phase 3 trials (NCT04646590) and has been approved for emergency use in Uzbekistan.

## Introduction

As of Feb 14, 2021, the coronavirus disease 2019 (COVID-19) pandemic caused by severe acute respiratory syndrome coronavirus 2 (SARS-CoV-2) infection (Coronaviridae Study Group of the International Committee on Taxonomy of, 2020; Jiang et al., 2020) has led to more than 100 million confirmed cases, with more than 2 million deaths (www.who.int). Vaccine is the “final solution” to control the pandemic. After 1 year of global efforts, a number of COVID-19 vaccines have been advanced into Phase 3 clinical trials (Dai and Gao, 2021). Several of them are approved for use in many countries, including mRNA vaccines, adenovirus vectored vaccines, inactivated vaccines and a protein subunit vaccine presented here (Anderson et al., 2020; Folegatti et al., 2020; Jackson et al., 2020; Logunov et al., 2020; Mulligan et al., 2020; Ramasamy et al., 2020; Sadoff et al., 2021; Sahin et al., 2020; Xia et al., 2020; Zhang et al., 2020; Zhu et al., 2020a; Zhu et al., 2020b). These approved vaccines mainly target whole virus or spike (S) protein (Dai and Gao, 2021). Different vaccines with different mechanisms of action would benefit for the cost-effective countermeasure to systemically stop the COVID-19 pandemic. Protein subunit represents as an important avenue to develop COVID-19 vaccines, with more than 80 vaccine candidates documented in World Health Organization (www.who.int). Currently, three protein subunit vaccines have been advanced in Phase 3 or 2/3 clinical trials, with two of them (NVX-CoV2327 and SCB-2019) using S protein as the antigen (Keech et al., 2020; Richmond et al., 2021).

In response to the pandemic, we developed a protein subunit vaccine targeting receptor-binding domain (RBD) (Dai et al., 2020; Yang et al., 2020). SARS-CoV-2 RBD located at C-terminal domain of S1 subunit in S protein, and is responsible for engagement of its cellular receptor human angiotensin-converting enzyme 2 (hACE2) (Wang et al., 2020). RBD is an attractive coronavirus vaccine target because it focuses the antibody response to blocking receptor binding, therefore, poses low potential for antibody-dependent enhancement (ADE) risk (Dai and Gao, 2021; Dai et al., 2020; Walls et al., 2020). To increase the immunogenicity, we designed a tandem-repeat dimeric RBD as the antigen for COVID-19 vaccine (Dai et al., 2020). Compared to the traditional monomeric RBD, RBD-dimer significantly enhanced the SARS-CoV-2 neutralizing antibodies produced in mice (Dai et al., 2020). We produced the RBD-dimer in Chinese hamster ovary (CHO) cell system, formulated with aluminum hydroxide as adjuvant. The resulting vaccine, ZF2001, is being evaluated in Phase 1 (NCT04445194, NCT04550351) and 2 (NCT04466085) clinical trials in China, showing good safety and immunogenicity (Yang et al., 2020). ZF2001 is now evaluated in international multi-center Phase 3 clinical trials (NCT04646590) and approved for emergency use in Uzbekistan. Here, we report the preclinical studies of ZF2001 in both mouse and non-human primate (NHP) models.

## Results

### Tandem-repeat RBD-dimer protein in ZF2001 vaccine

The RBD construct for SARS-CoV-2 started at S protein residue R319 and stopped at residue K537. Two copies of RBD were connected as tandem repeat dimer (Figure 1A). The construct of RBD-dimer was transformed into clinical-grade CHO cell lines. Cell lines with highest antigen yields were selected for scaling-up antigen production in accordance with current Good Manufacturing Practice. The tandem-repeat RBD-dimer antigen was further purified and characterized (Dai et al., 2020). The CHO-generated RBD-dimer bound to hACE2 with affinity similar as RBD-monomer, indicating the correctly exposure of receptor-binding motif (Figure S1). Stock solution was further formulated with aluminum hydroxide as adjuvant and put into vials as ZF2001 vaccine.

**Figure 1:**
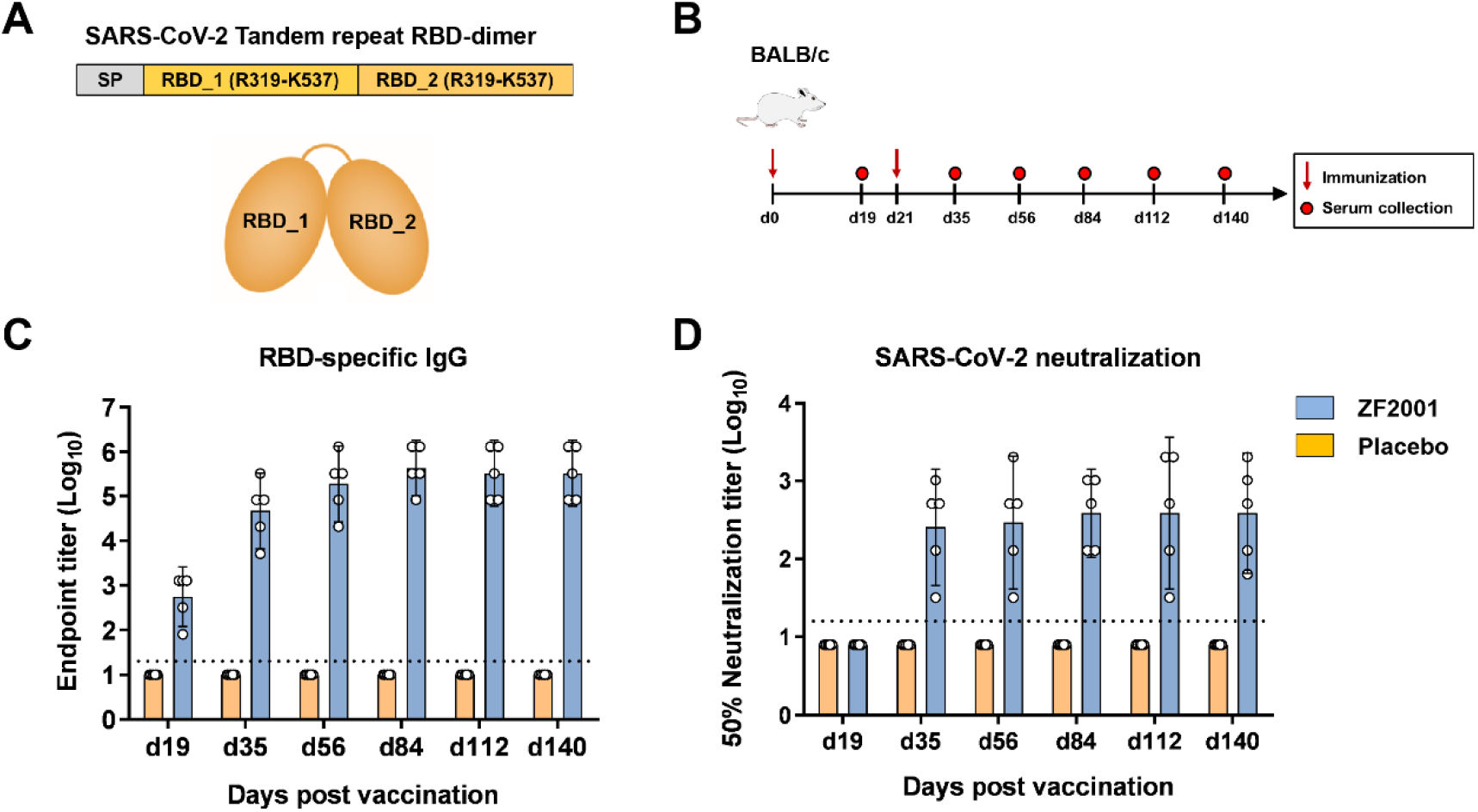
Humoral immune responses to ZF2001 vaccination in BALB/c mice. (A) schematic diagram of SARS-CoV-2 RBD dimer protein. Two copies of RBD from R319 to K537 are connected as tandem-repeat dimer. SP, signal peptide. (B) Time courses of ZF2001 vaccine immunization and sampling in groups of 6-8-week-old female BALB/c mice (n=5) vaccinated with 10 μg ZF2001 or placebo. (C) Enzyme-linked immunosorbent assay (ELISA) show serum IgG against SARS-CoV-2 RBD. The dashed line indicates the limit of detection. Data are geometric mean with 95% CI. (D) SARS-CoV-2 (HB01 strain) neutralization assay shows the 50% neutralization titer. The dashed line indicates the limit of detection. Data are geometric mean with 95% CI.

### ZF2001 vaccine immunogenicity in mice

To study the vaccine immunogenicity, groups of BALB/c mice were vaccinated with two doses of 10 μg ZF2001, 21 days apart. Mice receiving placebo (adjuvant-only) were used as the negative control. Serum samples were collected at different time points post vaccination to monitor the duration of antibody responses (Figure 1B). One immunization of ZF2001 vaccine induced a geometric mean titer (GMT) of 557 for serological RBD-binding IgG (Day 19), and this titer was further increased to 47,051 after a second immunization (Day 35) (Figure 1C). For vaccine-induced SARS-CoV-2 neutralizing antibodies (NAbs), the GMT was 256 after two immunizations at Day 35 (Figure 1D). These titers remain high for each time point until our latest sampling at Day 140, with GMTs of RBD-binding IgG from 188,203 to 432,376 (Figure 1C), and GMTs of SARS-CoV-2 NAb from 294 to 388 (Figure 1D), suggesting the durable humoral responses induced by ZF2001.

### ZF2001-elicited protection in Ad5-hACE2 transduced mice

To study the protective efficacy of ZF2001 vaccine against SARS-CoV-2 infection, groups of C57/B6 mice were immunized with two doses of 10 μg ZF2001, 21 days apart (Figure 2A). Mice receiving placebo were used as the control. Two doses of ZF2001 vaccine elicited high levels of both RBD-binding IgG (GMT, 81,920) and SARS-CoV-2 NAb (GMT, 959) (Figure 2B and 2C). These immunized mice were subsequently transduced via intranasal route with adenovirus expressing hACE2 as the SARS-CoV-2-sensitive animal model. Five days later, these transduced mice were intranasally challenged with 5 × 10^5^ or 1 × 10^5^ 50% tissue culture infectious dose (TCID_50_) of SARS-CoV-2 (HB01 strain) (Wei et al., 2020; Tan et al., 2020). Mice were euthanized and necropsied at 3 or 5 days post infection (DPI) to detect viral loads and exam pulmonary pathology. Encouragingly, in mice challenged with 5 × 10^5^ TCID_50_ SARS-CoV-2, the mean titers of viral genomic RNA (gRNA) per gram of lung for placebo and ZF2001 group were 6.5 × 10^8^ and 3.0 × 10^6^ (218-fold reduction), respectively, at 3 DPI, and 7.0 × 10^8^ and 4.8 × 10^5^ (1,448-fold reduction), respectively, at 5 DPI (Figure 2D). In mice challenged with 1 × 10^5^ TCID_50_ SARS-CoV-2, the mean gRNA titers for placebo and ZF2001 group, were 2.3 × 10^9^ and 1.2 × 10^5^, respectively, a reduction of 18,736-fold. The residual viral RNA in lung were probably derived from the high amount of input viruses. Therefore, we measured the magnitudes of subgenomic RNA (sgRNA) to quantify the infectious virus because sgRNA are generated in the infected cells during virus replication but is absent in virions (Kim et al., 2020). In mice challenged with 5 × 10^5^ TCID_50_ SARS-CoV-2, the mean titers of viral sgRNA per gram of lung for placebo recipients were 3.6 × 10^8^ and 5.1 × 10^8^ at 3 DPI and 5 DPI, respectively. By contrast, this titer was reduced to 1.4 × 10^6^ at 3 DPI and undetectable at 5 DPI for ZF2001 vaccine recipients (Figure 3B). Accordingly, in mice challenged with 1 × 10^5^ TCID_50_ SARS-CoV-2, these sgRNA titer was 8.1 × 10^8^ for placebo recipients, but was undetectable for ZF2001 recipients, at 3 DPI (Figure 2E). The neutralizing titers are negatively correlated with the gRNA and sgRNA, with Spearman correlation r value of -0.7127 and -0.6982, respectively (Figure 2F). Consistent with this, immunofluorescence analysis of lung section stained with anti-SARS-CoV-2 NP antibody demonstrated the presence of viral protein in mice receiving placebo, but absence in mice receiving ZF2001 vaccine (Figure 3A). These results demonstrated that ZF2001 vaccine is protective against pulmonary SARS-CoV-2 infection.

**Figure 2:**
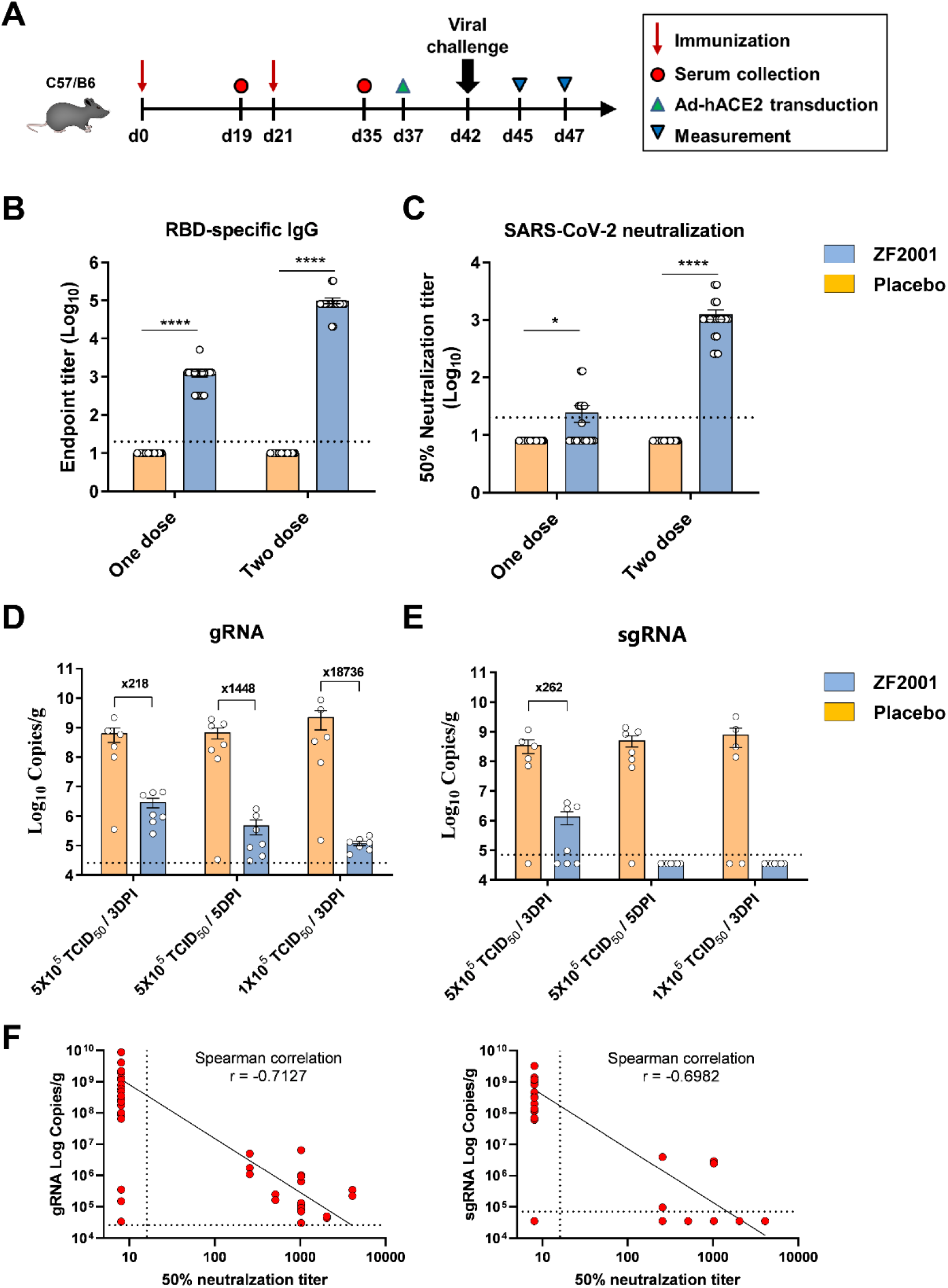
Protective efficacy of ZF2001 vaccine in mice. (A) Time course of immunization, sampling, viral challenge and measurement. Two groups of C57/B6 mice (n=18) were immunized with two doses of 10 μg ZF2001 vaccine or placebo, 3 weeks apart. Serum samples were collected after both one and two doses as indicated. Mice were then transduced with 8 × 10^8^ vp of Ad5-hACE2 via intranasal (i.n.) route. Each groups of mice were further split into three groups (n=6), with the former two groups infected with high-dose (5 × 10^5^ TCID_50_) SARS-CoV-2 and the latter group infected with low-dose (1 × 10^5^ TCID_50_) SARS-CoV-2. Lung tissues were harvested at either 3 or 5 DPI for the two groups with high-dose virus challenge and 3 DPI for the group with the low-dose virus challenge. (B) ELISA shows serum IgG against SARS-CoV-2 RBD. Data are geometric mean with 95% CI. P values were analyzed with unpaired t test (****, P < 0.0001). The dashed line indicates the limit of detection. (C) SARS-CoV-2 (HB01 strain) neutralization assay shows the 50% neutralization titer. Data are geometric mean with 95% CI. P values were analyzed with unpaired t test (*, P < 0.05; ****, P < 0.0001). The dashed line indicates the limit of detection. (D-E) SARS-CoV-2 titration from lung tissues by qRT-PCR probing virus Grna and sgRNA (E). Data are means ± SEM. The dashed lines indicate the limit of detection. (F) Protective correlation of NAb titer with SARS-CoV-2 gRNA or sgRNA, calculated with Spearman correlation in GraphPad Prism 9.0.

**Figure 3:**
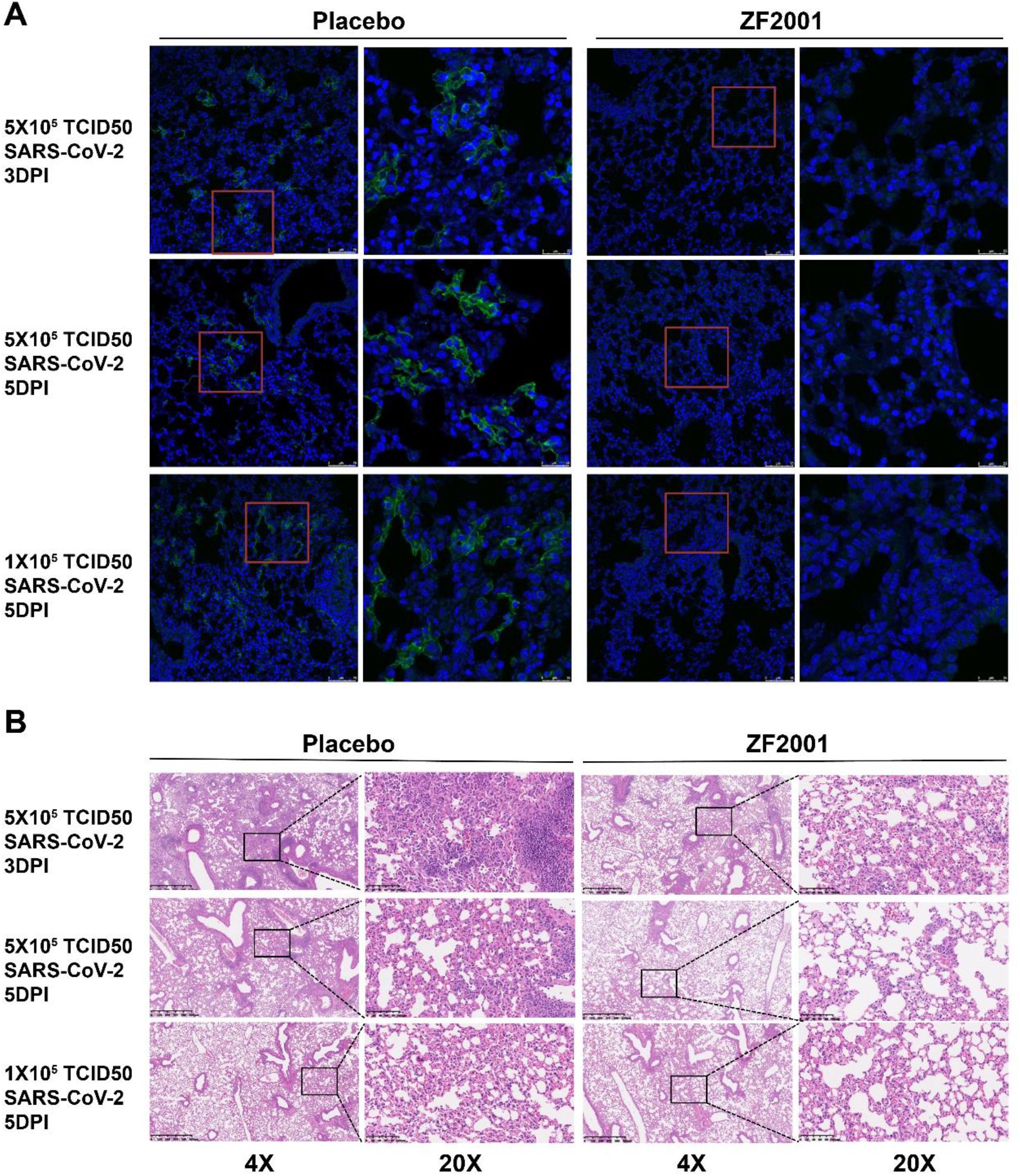
Immunofluorescence and histopathology analyses of lung tissues in mice. (A) Immunofluorescence analysis of lung tissue section stained with anti-SARS-CoV-2 nucleoprotein (N) antibody. Green: SARS-CoV-2 N protein; Blue: DAPI. (B) Typical histopathology images of lung tissues section shown by hematoxylin-eosin staining. Both low magnifications and high magnifications are shown, highlighted by boxes.

To further assess the vaccine protection against SARS-CoV-2-induced lung damage, histopathological examination was performed. All lung tissue samples harvested from mice with placebo exhibited apparently moderated to severe viral pneumonia, characterized by thickened alveolar walls, vanishment of alveolar cavities, pulmonary vascular congestion and diffuse inflammatory cell infiltration (Figure 3B). By contrast, a markedly relieved histopathological changes were observed in lung tissues of mice vaccinated with ZF2001 vaccine (Figure 3B). This result demonstrated that ZF2001 protected against SARS-CoV-2-induced lung injury in mice, without observation of vaccine enhanced diseases.

### ZF2001 vaccine immunogenicity in cynomolgus macaques

To further evaluate ZF2001 vaccine in NHPs, groups of healthy young cynomolgus macaques (n=10) were immunized with four doses of 25 μg or 50 μg vaccine intramuscularly at Week 0, 4, 8 and 10, respectively, to monitor the immunogenicity kinetics (Figure 4A). We found after one immunization, ZF2001 vaccine can elicit serological RBD-binding IgG GMTs up to ∼100,000 in both the 25 μg or 50 μg groups, and these titers were further enhanced to more than 1000,000 after the second immunization (Figure 4B). The third and fourth boosts did not significantly increase the GMTs (Figure 4B). For the NAb response, two immunizations of ZF2001 vacine elicited SARS-CoV-2 neutralizing GMTs of 630 in the 25 μg group and 776 in the 50 μg group, respectively. The third immunization did not further enhance NAb titer (Figure 4C).

**Figure 4:**
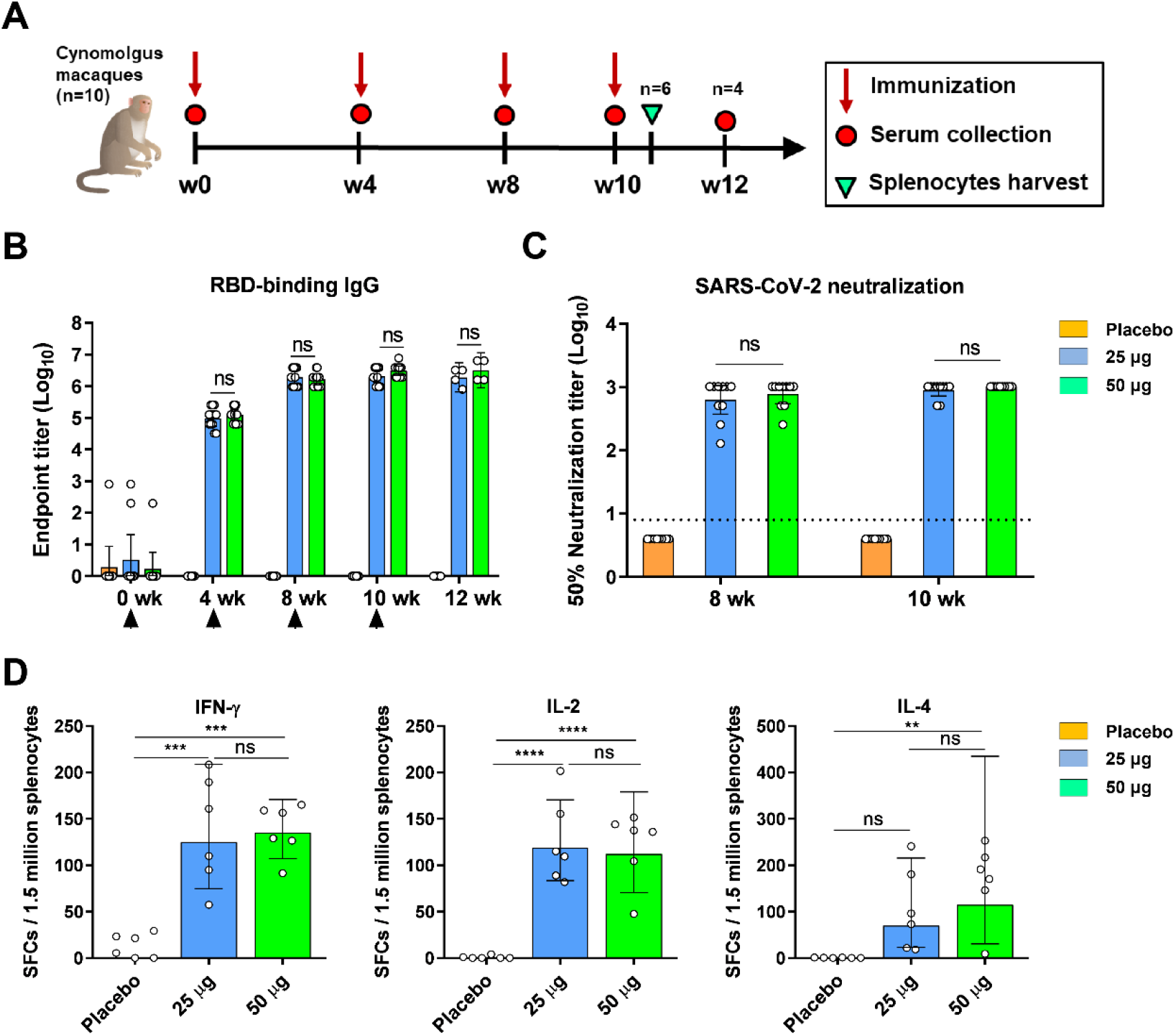
Humoral and cellular immune responses to ZF2001 vaccination in cynomolgus macaques. (A) Time course of immunization and sampling. Groups of cynomolgus macaques (n=10) were immunized with four doses of placebo, the 25 μg vaccine and the 50 μg vaccine, respectively. Serum samples were collected at indicated time points post vaccination. Six macaques in each group were euthanized for spleen harvest at day 3 after the last vaccination. (B) ELISA shows serum IgG against SARS-CoV-2 RBD. Data are geometric mean with 95% CI. P values were analyzed with unpaired t test (ns, not significant). (C) SARS-CoV-2 (HB01 strain) neutralization assay shows the 50% neutralization titer. Data are geometric mean with 95% CI. P values were analyzed with unpaired t test (ns, not significant). The dashed line indicates the limit of detection. (D) Splenic IFN-γ, IL-2 and IL-4 ELISPOT responses to SARS-CoV-2 RBD antigen. SFCs: Spot-forming cells. Data are geometric mean with 95% CI. P values were analyzed with one-way ANOVA (ns, not significant; **, P < 0.01; ***, P < 0.001; ****, P < 0.0001)

To detect the cellular immune responses in NHPs, 6 macaques in each group were euthanized to harvest spleen at 3 days after the last vaccination. Splenocytes were stimulated with RBD protein. ELISPOT assay was perform to detect T_H_1 (IFN-γ, IL-2) and T_H_2 (IL-4) cytokine production. We found both 25 μg or 50 μg ZF2001 induced substantial cellular responses, with the significantly enhanced and T_H_1/T_H_2 balanced cytokine production (Figure 4D).

Overall, macaques receiving 50 μg vaccine did not show enhanced immunogenicity compared to those receiving 25 μg vaccine for both humoral and cellular immune responses.

### ZF2001-elicited protection in rhesus macaques

To assess the protection efficacy, we immunized healthy young rhesus macaques (n=3) with two doses of 25 μg or 50 μg vaccine intramuscularly, 21 days apart (Figure 5A). Macaques receiving placebo was used as the control. Both 25 μg and 50 μg ZF2001 vaccine elicited high levels of serological RBD-binding IgG and SARS-CoV-2 neutralizing antibody after one or two immunizations. For RBD-binding IgG, the GMTs reached 6,451 for both the 25 μg and 50 μg groups after the first immunization (Figure 5B). After the second immunization, the GMTs further raised to 100,794 and 80,000 for the 25 μg and 50 μg group, respectively, at Day 28 (Figure 5B). No significant increase was observed at Day 35, with the GMTs of 63,496 for the 25 μg group and 50,397 for the 50 μg group (Figure 5B). For SARS-CoV-2 NAbs, two immunizations of ZF2001 vaccine elicited GMTs of 256 for the 25 μg group and 161 for the 50 μg group at Day 28 (Figure 7C). No significant increase was observed at Day 35, with the GMTs of 203 for the 25 μg group and 256 for the 50 μg group (Figure 5C). The 50 μg group did not show enhanced immunogenicity compared to the 25 μg group in macaques.

**Figure 5:**
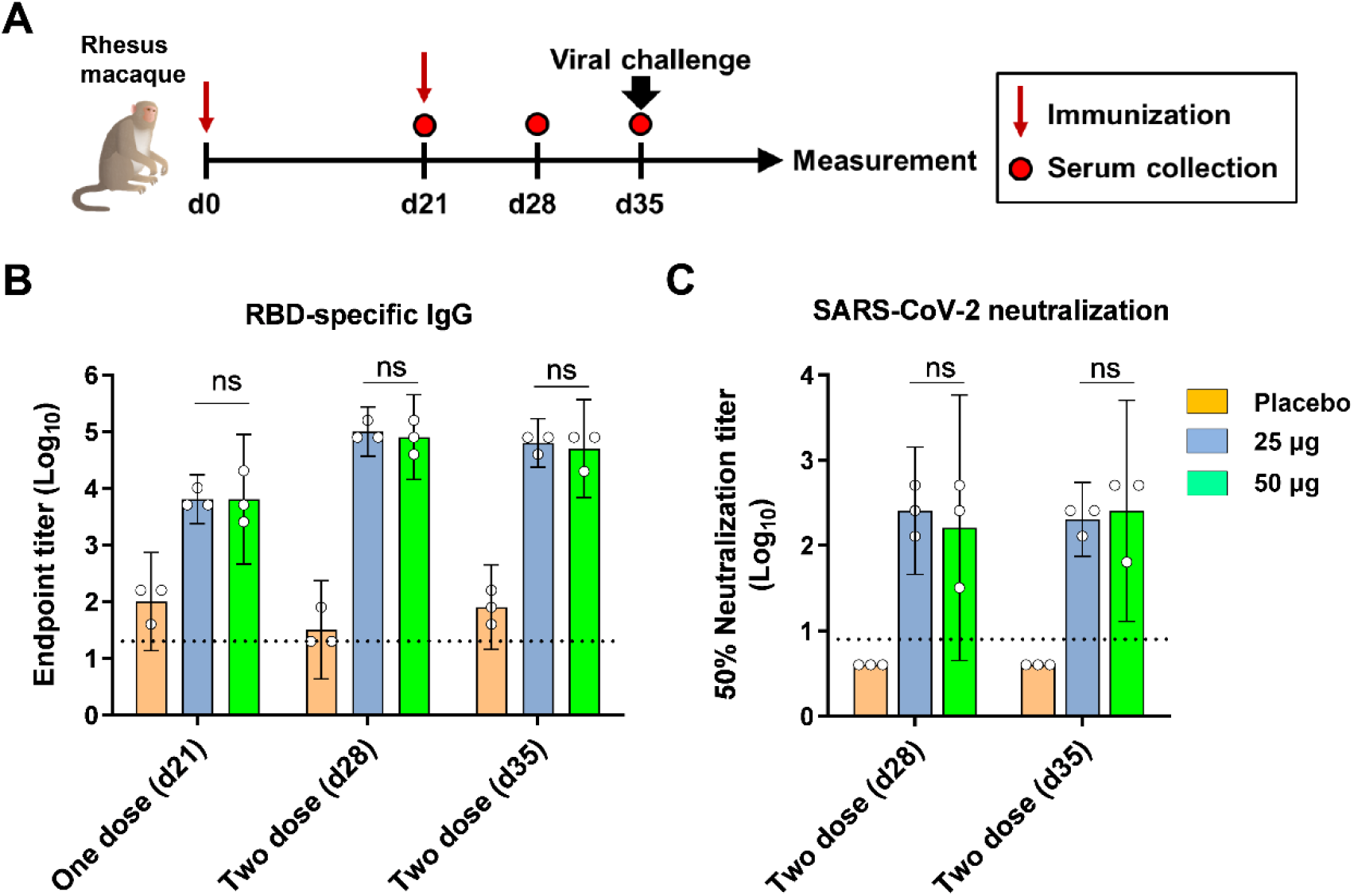
Humoral immune responses to ZF2001 vaccination in rhesus macaques. (A) Time course of immunization, sampling, viral challenge and measurement. Groups of rhesus macaques (n=3) were immunized with two doses of placebo, the 25 μg vaccine and the 50 μg vaccine. Serum samples were collected at indicated time points post vaccination. At 14 post the second immunization, macaques were challenged with 2 × 10^6^ TCID_50_ SARS-CoV-2 (20SF107 strain) via intratracheal route. Macaques were euthanized at 7 DPI for tissue harvest. (B) ELISA shows serum IgG against SARS-CoV-2 RBD. Data are geometric mean with 95% CI. P values were analyzed with unpaired t test (ns, not significant). (C) SARS-CoV-2 neutralization assay shows the 50% neutralization titer. Data are geometric mean with 95% CI. P values were analyzed with unpaired t test (ns, not significant). The dashed line indicates the limit of detection.

The macaques were challenged with 1 × 10^6^ TCID_50_ SARS-CoV-2 intratracheally at Day 28 post vaccination. Macaques were euthanized at 8 DPI. Tissues from 7 lung lobes (4 sites in each lobe), trachea, left and right bronchus were collected and quantified for viral gRNA. As expected, substantial virus loads can be detected in lung lobes (average 4748 copies/μg gRNA). By contrast, animals receiving either 25 μg or 50 μg ZF2001 vaccine exhibited significantly reduced virus loads in lung lobes, with average 1 and 10 copies/μg gRNA for 25 μg and 50 μg group, respectively. In addition, viral gRNA was detected in trachea and bronchi from 1 or 2 macaques receiving placebo, but was undetectable in all ZF2001-vaccinated macaques (Figure 6A).

**Figure 6:**
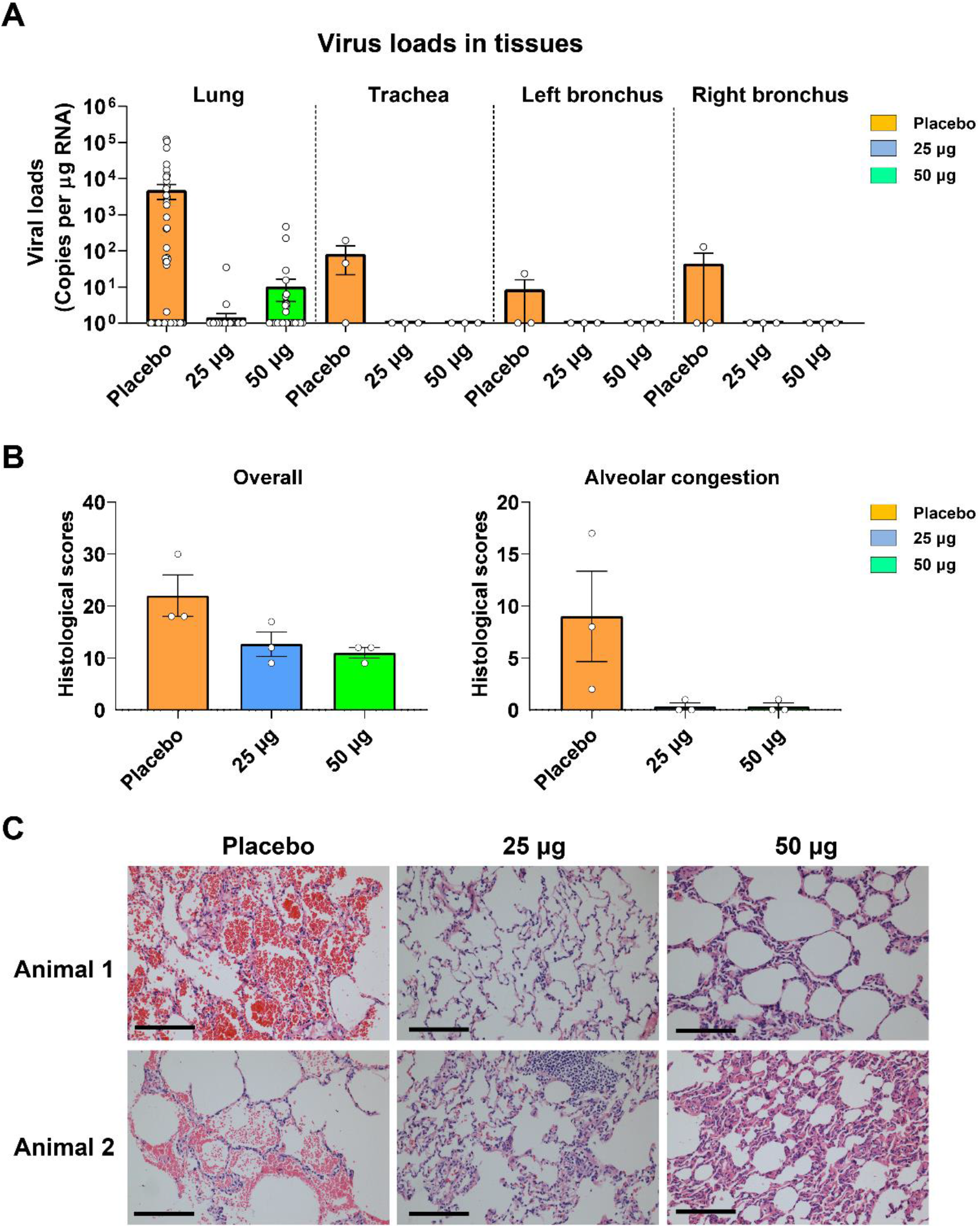
Viral loads and clinical signs in rhesus macaques challenged with SARS-CoV-2 after vaccination. (A) SARS-CoV-2 titration from lung tissues by qRT-PCR probing virus gRNA. Tissue samples: 7 lung lobes (4 sites for each lobes) of three macaques in each group (84 samples per group); tracheas of three macaques in each group; left and right bronchi of three macaques in each group. (B) Histopathology scores of overall lung lesions and pulmonary alveolar congestion. (C) Typical histopathology images of lung tissues section shown by hematoxylin-eosin staining. Two macaques from each group. Scale bar, 100 μm.

To further assess the protection efficacy in lung, pulmonary histopathology of each macaque was scored based on the thickening of alveolar septa, pulmonary alveolar congestion and inflammatory cell infiltration in alveoli and trachea. Overall, high scores of lung lesions were found in control animals, but were dramatically reduced in ZF2001-vaccinated animals (Figure 6B). In particular, all three control animals showed severe pulmonary alveolar congestion. By contrast, macaques receiving either 25 μg or 50 μg vaccine showed almost no alveolar congestion (Figure 6B and 6C). The typical images of tissue hematoxylin-eosin (H/E) staining were shown in the Figure 6C. We did not observe vaccine enhanced diseases in all these macaques.

## Discussion

ZF2001 is a tandem-repeat dimeric RBD protein COVID-19 vaccine currently under Phase 3 clinical trials (NCT04646590) and emergency use in Uzbekistan. Here, we report its preclinical results in animal models of both mice and NHP. ZF2001 vaccine was highly immunogenic in both mouse and NHP models, with high neutralizing GMTs against SARS-CoV-2. Two shots of vaccine protected both hACE2-transduced mice and NHPs against SARS-CoV-2 infection. No evidence of vaccine enhanced diseases was found in both models. Compared with the other two protein subunit vaccines advanced in Phase 3 or 2/3 clinical trials (NVX-CoV2373 and SCB-2019) using full-length S protein as antigen, ZF2001 vaccine is an RBD-based protein vaccine aiming to focus immune responses in receptor-binding blocking, with a unique design of dimer for better stability and immunogenicity (Dai et al., 2020). Besides, ZF2001 vaccine uses a traditional alum-based adjuvant with long safety profiles in humans, rather than the relatively new adjuvants used in NVX-CoV2373 (Matrix-M) and SCB-2019 (AS03 or CpG1018+Alum) (Keech et al., 2020; Richmond et al., 2021). Besides, ZF2001 is stable in 2-8 °C and allows scalable manufacturing due to its high antigen yields (Dai et al., 2020). This highlights its advantage of vaccine deployment to meet the global vaccine demands. Therefore, the encouraging preclinical results suggest that ZF2001 vaccine with different vaccine target, different adjuvant and different mechanism of action would diversify the current vaccine pipelines.

In NHPs models, the higher vaccine dose (50 μg) did not show a superior immune response or better protection compared to the lower vaccine dose (25 μg). We speculate that 25 μg dose has already reached or is near to a saturated dosage to stimulate immune system. Larger dose may not further increase the immunogenicity. On the contrary, sometimes larger dose may decrease the protection as reported for an adenovirus-based vaccine AZD1222 (Voysey et al., 2021). The similar immunological trend was also observed in Phase 1 and 2 clinical trials (Yang et al., 2020). This preclinical results in NHPs support the use of 25 μg vaccine dose to an ongoing Phase 3 large scale evaluation for safety and efficacy.

## Author contributions

L.D., J.Y., and G.F.G. initiated and coordinated the project. Y.A., J.Y., L.D., and G.F.G. designed the experiments. Y.A., S.L., X.J., J.H., S.X., K.X., Y.H., M.L., M.Y.,C.L., T.Z., T.S., B.H., L.Z., W.W., R.A., Y.C., C.W., E.H., S.Y., Y.B., and C.K. conducted the experiments. G.W. produced the Ad5-hACE2 virus. J.H., and T.S. performed the monkey experiment and Y.Z supervised it. Y.A., S.L. X.J. and K.X. performed the mice challenge experiment and conducted the live virus neutralization assay for SARS-CoV-2 vaccine. Y.A., S.L., J.H., T.Z., L.D. and G.F.G. analyzed the data. L.D., and Y.A., wrote the manuscript. L.D., Y.A., K.X., and G.F.G. discussed and edited the manuscript.

## Acknowledgements

This work is supported by the Strategic Priority Research Program of the Chinese Academy of Sciences, China (XDB29010202), the National Program on Key Research Project of China (2020YFC0842300 and 2020YFA0907102), the National Natural Science Foundation of China, China (NSFC) (81991494 and 82041048), the intramural special grant for SARS-CoV-2 research from the Chinese Academy of Sciences and Anhui Zhifei Longcom Biopharmaceutical. Lianpan Dai is supported by Youth Innovation Promotion Association CAS, China (2018113).

## Declaration of interests

Y.A., K.X., X.H., T.Z., J.Y., L.D. and G.F.G. are listed in the patent as the inventors of the RBD-dimer as a betacoronavirus vaccine. All other authors declare no competing interests.

**Figure S1:**
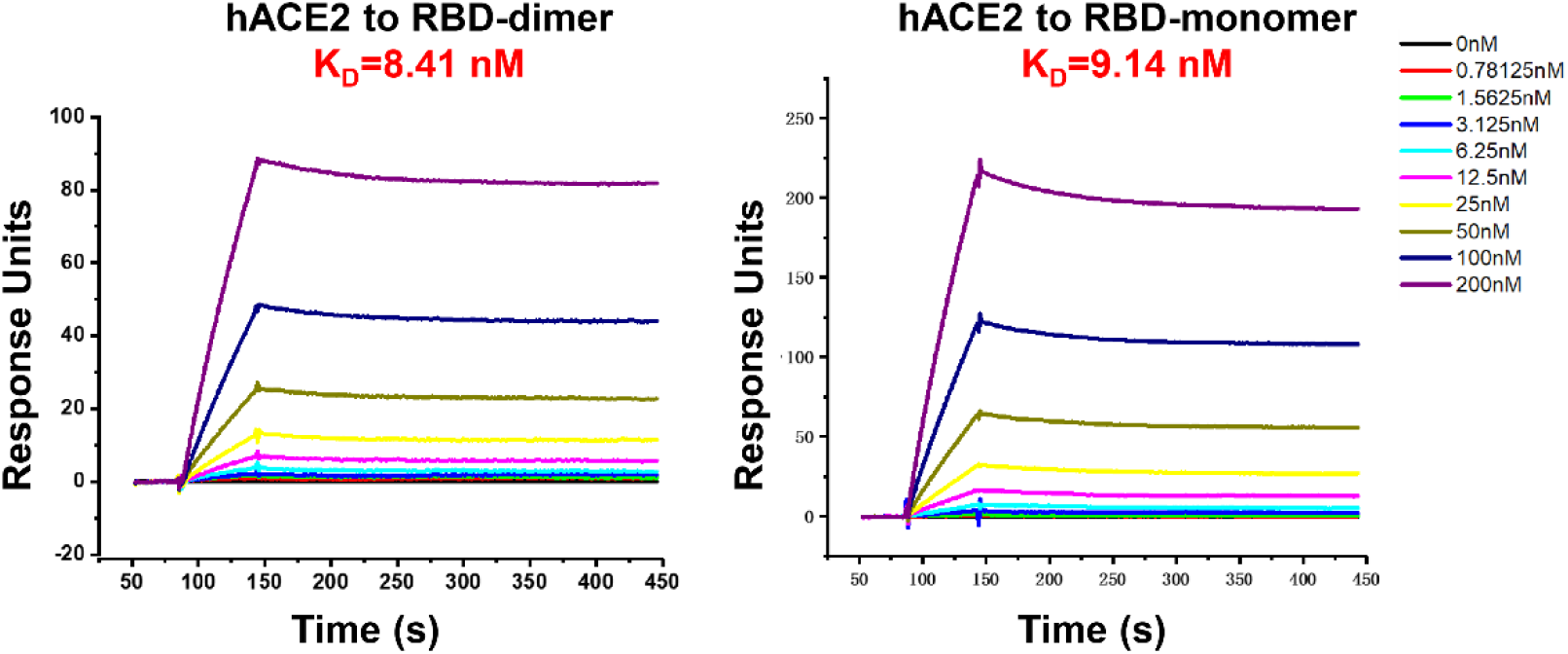
Representative BIAcore diagrams of RBD-dimer and RBD-monomer bound to hACE2 protein. The K_D_ value was calculated by the software BIAevaluation Version 4.1 (GE Healthcare).

## Supplementary

### Materials and Methods

#### Mouse experiments

Female BALB/c mice and female C57BL/6 mice were immunized intramuscularly (i.m) with ZF2001 (10 μg dose antigen) or placebo and boosted with equal dose of vaccine at day 21 post priming. Serum samples were collected after vaccination as indicated in figures legends.

For challenge experiment, C57BL/6 mice were intranasally (i.n.) transduced with 8 × 10^8^ vp of Ad5-hACE2 as a mouse model for SARS-CoV-2 infection (Hassan et al., 2020). Five days later, the transduced mice were infected with 5 × 10^5^ or 1 × 10^5^ TCID_50_ of SARS-CoV-2 (HB01 strain) via i.n. route. The mice were euthanized and necropsied 3 days after challenge. Lung tissues were collected for virus titration and pathological examination. All mice experiments with SARS-CoV-2 challenge were conducted under animal biosafety level 3 (ABSL3) in IMCAS.

#### Monkey experiments

Cynomolgus macaques were divided into three groups (n=10), with five females weighing 2.97 to 3.62 kg and five males weighing 2.95 to 3.67 kg in each group. Macaques were immunized with 25 μg or 50 μg ZF2001 vaccine via i.m. route at Week 0, 4, 8 and 10. Group receiving placebo was used as the control. Serum samples were collected for detection of SARS-CoV-2-specific IgG and neutralizing antibodies. Spleens were harvested to prepare splenocytes for detection of cytokines (IFN-γ, IL-2 and IL-4) by <u>ELISPOT Assay</u>.

Rhesus macaques at 3 to 6 years of age were divided into three groups (n=3), with one female and two male monkeys in each group. Macaques were immunized with two doses of 25 μg or 50 μg ZF2001 via i.m. route, 21 days apart. Serum samples were collected for detection of SARS-CoV-2-specific IgG and neutralizing antibodies.

For challenge experiment, rhesus macaques were infected with 2×10^6^TCID_50_ of SARS-CoV-2 (20SF107 strain) via intratracheal route. The experimental animals were anesthetized by injection of Zoletil 50 (Virbac, France) into the thigh muscle. Euthanasia was performed on Day 8 after the challenge. Tissues from lungs (7 lobes with 4 sites in each lobe), trachea, left and right bronchus were collected for viral genomic RNA quantification and histopathology staining.

#### Protein expression and purification

Monomeric RBD protein of SARS-CoV-2 used in ELISA assay was expressed and purified as previously described (Dai et al., 2020). Briefly, the coding sequence for SARS-CoV-2 RBD (S protein 319-541, GISAID accession No. EPI_ISL_402119) was codon-optimized for mammalian cell expression and synthesized. For this construct, signal peptide sequence of MERS-CoV S protein (S protein residues 1-17) was added to the protein N terminus for protein secretion, and a hexa-His tag was added to the C terminus to facilitate further purification processes. The construct, which synthesized by GENEWIZ, China, was cloned into the pCAGGS vector and transiently transfected into HEK293T cells. After 3 days, the supernatant was collected and soluble protein was purified by Ni affinity chromatography using a HisTrap ^™^ HP 5 mL column (GE Healthcare). The sample was further purified via gel filtration chromatography with HiLoad® 16/600 Superdex® 200 pg (GE Healthcare) in a buffer composed of 20 mM Tris-HCl (pH 8.0) and 150 mM NaCl.

#### ELISA

For mice and rhesus macaque, ELISA plates (3590; Corning, USA) were coated over-night with 3 μg/ml of RBD protein in 0.05 M carbonate-bicarbonate buffer, pH 9.6, and blocked in 5% skim milk in PBS. Serum samples were serially diluted and added to each well. Plates were incubated with goat anti-mouse IgG-HRP antibody or goat anti-monkey IgG-HRP antibody and subsequently developed with 3,3’,5,5’-tetramethylbenzidine (TMB) substrate. Reactions were stopped with 2 M hydrochloric acid, and the absorbance was measured at 450 nm using a microplate reader (PerkinElmer, USA). The endpoint titers were defined as the highest reciprocal dilution of serum to give an absorbance greater than 2.5-fold of the background values. Antibody titer below the limit of detection was determined as half the limit of detection.

For cynomolgus macaques, ELISA plates were coated over-night with 1 μg/ml of RBD-dimer protein and blocked in 3% skim milk in PBST. Serum samples were serially diluted and added to each well. 10ul serum from cynomolgus macaques of blank control group was mixed as one blank control serum and diluted as the same with initial fold of vaccine group. The subsequent operations were as described above. Cut off value was defined 2.1-fold of OD_450_ values of the blank control serum. When OD_450_ > Cut off value, the serum was determined positive. The endpoint titer was defined as the highest reciprocal dilution of serum with an absorbance greater than 2.1-fold of the blank control serum value. When OD_450_ value of the serum to test ≤ Cut off value, the endpoint titer was zero.

#### Live SARS-CoV-2 neutralization assay

The neutralizing activity of serum was assessed using a previously described SARS-CoV-2 neutralization assay (Nie et al., 2020). Briefly, mouse serum samples were 4-fold serially diluted and mixed with the same volume of 100 TCID_50_ SARS-CoV-2, then incubated at 37°C for 1 hour. Then, 100 μL virus-serum mixture was transferred to pre-plated Vero cells in 96-well plates. Inoculated plates were incubated at 37°C for an additional 72 hours to monitor the cytopathic effect (CPE) microscopically. The neutralization titers were defined as the reciprocal of serum dilution required for 50% neutralization of viral infection. All the live virus neutralization assay of mouse serum samples was conducted under biosafety level 3 (BSL3) facility in IMCAS, with SARS-CoV-2 HB01 strain. The neutralizing activity of serum samples from macaques were assessed with 2020XN4276 strain of SARS-CoV-2 in the same method as mentioned above in Guangdong Provincial Center for Disease Control and Prevention

#### ELISPOT

To detect antigen-specific T cell responses, ELISPOT assay was performed as previously described (Xu et al., 2018), with some modifications. Briefly, Flat-bottom, 96-well plates which were precoated with 10 μg/ml anti-mouse IFN-γ Ab (BD Biosciences, USA) overnight at 4°C, pre-coated plates (mAb MT126L or mAb 2A91/2C95) and flat-bottom, 96-well plates which were precoated with anti-monkey IL-4 Ab (U-Cytech) overnight at 4°C were washed with sterile PBS and followingly blocked according to guidelines for Mouse IFN-γ ELISPOT kit (BD), Monkey IFN-γ ELISPOT ^PLUS^ kit (MABTECH), IL-2 ELISPOT^PLUS^ kit (MABTECH) or IL-4 ELISPOT kit (U-Cytech). Splenocytes of mice or cynomolgus macaques were added to the plate. Peptide pool covering RBD of SARS-CoV-2 (2 μg/ml individual peptide) (for mouse) or RBD-dimer protein of SARS-CoV-2 (for cynomolgus macaques) was added to the wells for stimulation. Phytohemagglutinin (PHA) was added as a positive control. Cells without stimulation were employed as a negative control. After 12-36 hours of incubation, the cells were removed, and the plates were processed with biotinylated IFN-γ, IL-2 or IL-4 detection antibody, streptavidin-HRP conjugate, and substrate. When the colored spots were intense enough to be observed, the development was stopped by thoroughly rinsing samples with deionized water. The numbers of the spots were determined using an automatic ELISPOT reader and image analysis software.

#### qRT-PCR for mice

Mice lung tissues were weighed and homogenized. Virus genomic RNA was isolated from 50-μl supernatants of homogenized tissues using a nucleic acid extraction instrument MagMAX^™^ Express Magnetic Particle Processor (Applied Biosystems, USA). SARS-CoV-2-specific quantitative reverse transcription-PCR (qRT-PCR) assays were performed using a FastKing One Step Probe RT-qPCR kit (Tiangen Biotech, China) on a CFX96 Touch real-time PCR detection system (Bio-Rad, USA) according to the manufacturer’s protocol. Two sets of primers and probes were used to detect a region of the N gene of viral genome (Chandrashekar et al., 2020) and a region of E gene of subgenomic RNA (sgRNA) from SARS-CoV-2 (Wolfel et al., 2020) respectively, with sequences as follows:

N-gene-F, GACCCCAAAATCAGCGAAAT;

N-gene-R, TCTGGTTACTGCCAGTTGAATCTG;

N-gene-probe, FAM-ACCCCGCATTACGTTTGGTGGACC-TAMRA (where FAM is 6-carboxyfluorescein, and TAMRA is 6-carboxytetramethylrhodamine);

sgRNA-E-F, CGATCTCTTGTAGATCTGTTCTC;

sgRNA-E-R, ATATTGCAGCAGTACGCACACA;

sgRNA-E-probe, FAM-ACACTAGCCATCCTTACTGCGCTTCG-TAMRA.

Viral loads were expressed on a log10 scale as viral copies/gram after calculation with a standard curve. Viral copy numbers below the limit of detection were set as the half of the limit of detection.

#### qRT-PCR for rhesus macaques

The total RNA of tissues from rhesus macaques were extracted with TRIzol reagent method (Thermo USA) (Song et al., 2020), Viral RNA detection were detected with a probe one-step real-time quantitative PCR kit (TOYOBO, Japan). Previously reported primers (5’-GGGGAACTTCTCCTGCTAGAAT-3’, 5’-CAGACATTTTGCTCTCAAGCTG-3’) targeting SARS-CoV-2 N protein and probe (FAM-TTGCTGCTGCTTGACAGATT-TAMRA-3’) were used (Tian et al., 2021). The dilution of each test run was referred to the standard (National Institute of Metrology, China), and the copy number of each sample was calculated.

#### Histopathology analysis

Mice lung tissues were fixed in 4% paraformaldehyde, dehydrated, embedded in paraffin, and then sectioned. Tissue sections (4 μm) were deparaffinized in xylene and stained with hematoxylin and eosin (H&E) for pathological examination, such as peribronchiolitis, interstitial pneumonitis and alveolitis. All the rhesus macaque tissues were fixed in 4% paraformaldehyde for minimum of seven days, then embedded in paraffin, sectioned (slices 4 microns) for HE staining (Zhang et al., 2016).

Pulmonary histopathology of lung tissue was scored based on the thickening of alveolar septa, pulmonary alveolar congestion and inflammatory cell infiltration in alveoli and trachea. For thickening of alveola septa, no significant widened alveolar septa was observed is scored 0; slightly widened alveolar septa with lesion range < 25% is scored 1; moderate widened, fused and consolidated alveolar septa with lesion range 25%-50% is scored 2; severe widened, fused, consolidated alveolar septa with lesion range 50%-75% is scored 3; extremely severe widened, fused, and consolidated alveolar septa with lesion range >75% is scored 4. For inflammatory cell infiltration and alveolar congestion, no lesion or disease variable is scored 0; mild/small lesions with lesion range or disease variable < 25% is scored 1; moderate lesions with lesion range or disease variable 25%-50% is scored 2; severe/multiple lesions with lesion range or disease variable 50%-75% is scored 3; extremely severe/numerous lesions with lesion range or disease variable 50%-75% is scored 4.

#### Immunofluorescence

The paraffin tissue sections were deparaffinized with xylene, rehydrated through successive bathes of water and incubated in 3% H_2_O_2_ at room temperature. Subsequently, the sections were blocked with BSA, incubated with primary antibody against SARS-CoV-2 nucleoprotein (Sino Biological) at 37 °C and then incubated with goat anti-rabbit IgG (H+L) Alexa Fluor 488 antibody (Abways). DAPI (4’,6-diamidino-2-phenylindole) was also incubated with the sections, followed by detection using laser scanning confocal microscope (Leica).

#### SPR

Protein interactions were tested through SPR analysis and the experiments were carried out at 25°C using a BIAcore 3000 machine with CM5 chips (GE Healthcare). All proteins for SPR analysis were exchanged to PBST (10 mM Na2HPO4; 2mM KH2PO4, pH 7.4; 137 mM NaCl; 2.7 mM KCl; 0.005% Tween 20). SARS-CoV-2 RBD proteins were immobilized onto CM5 chips and analyzed for real-time binding by flowing through gradient concentrations of hACE2. The aforementioned RBD proteins were immobilized on the chip at about 1000 response units (RUs). The concentrations of hACE2 are 0,0.78125,1.5625,3.125,6.25,12.5,25,50,100, and 200nM. After each cycle, the sensor surface was regenerated using 7 μl of 10 mM NaOH. Measurements from the reference flow cell (immobilized with BSA) were subtracted from experimental values. The binding kinetics were analyzed using 1:1 binding model with the software BIAevaluation Version 4.1 (GE Healthcare).

### Statistics

Data was expressed as the means ± standard errors of the means (SEM). For all analyses, *p* values were analyzed with unpaired t test or one-way ANOVA. Correlation analysis was conducted with Spearman correlation.

